# Are Genetic Interactions Influencing Gene Expression Evidence for Biological Epistasis or Statistical Artifacts?

**DOI:** 10.1101/020479

**Authors:** Alexandra E. Fish, John A. Capra, William S. Bush

**Affiliations:** Vanderbilt Genetics Institute, Vanderbilt University, Nashville, Tennessee, 37235, USA; Department of Biological Sciences, Vanderbilt University, Nashville, Tennessee, 37235, USA; Department of Biomedical Informatics, Vanderbilt University, Nashville, Tennessee, 37235, USA; Institute for Computational Biology, Department of Epidemiology and Biostatistics, Case Western Reserve University, Cleveland, Ohio, 44106, USA

## Abstract

The importance of epistasis – or statistical interactions between genetic variants – to the development of complex disease in humans has long been controversial. Genome-wide association studies of statistical interactions influencing human traits have recently become computationally feasible and have identified many putative interactions. However, several factors that are difficult to address confound the statistical models used to detect interactions and make it unclear whether statistical interactions are evidence for true molecular epistasis. In this study, we investigate whether there is evidence for epistasis regulating gene expression after accounting for technical, statistical, and biological confounding factors that affect interaction studies. We identified 1,119 (FDR=5%) interactions within *cis*-regulatory regions that regulate gene expression in human lymphoblastoid cell lines, a tightly controlled, largely genetically determined phenotype. Approximately half of these interactions replicated in an independent dataset (363 of 803 tested). We then performed an exhaustive analysis of both known and novel confounders, including ceiling/floor effects, missing genotype combinations, haplotype effects, single variants tagged through linkage disequilibrium, and population stratification. Every replicated interaction could be explained by at least one of these confounders, and replication in independent datasets did not protect against this issue. Assuming the confounding factors provide a more parsimonious explanation for each interaction, we find it unlikely that *cis*-regulatory interactions contribute strongly to human gene expression. As this calls into question the relevance of interactions for other human phenotypes, the analytic framework used here will be useful for protecting future studies of epistasis against confounding.

## Introduction

Epistasis, a phenomenon wherein the effect of a genetic variant on the phenotype is dependent on other genetic variants, was first identified over a century ago; however, it has been highly contested whether or not epistasis plays an important role in the development of complex disease in humans. In model organisms, epistasis is commonly observed: variants associated with the trait of interest often interact with other variants, and more broadly, such interactions account for^1–3^ a notable proportion of variance in an array of phenotypes. Epistasis may play a similar role in humans as additive genetic effects are unable to account for the majority of heritability in most complex traits;^4,5^ however, evidence for epistasis in human remains elusive. Most studies rely on the statistical association between genetic variants and phenotype to identify signs of epistasis, and the interactions identified are notoriously difficult to replicate.^6,7^ This may be attributable to the inherent inability to tightly control a variety of factors when studying phenotypes in humans, or the fact that most phenotypes studied are several steps removed from the underlying biological processes. These methodological limitations make it unclear whether the lack of observed epistasis in humans is a true feature of the genetic architecture, or is simply much more difficult to observe outside experimental systems.

Human-derived cell lines, while a proxy for primary tissue, provide a unique opportunity to investigate epistasis. Like model systems, the environment can be tightly controlled, and moreover, comprehensive genetic and gene expression data can readily be collected from cell lines. Through statistical association studies, the genetic architecture underlying thousands of genes’ expression – a quantitative phenotype directly tied to the nucleotide sequence – can be interrogated. Furthermore, molecular mechanisms that drive gene expression are known to involve complex interactions among transcription factors and regulatory sequences, and experimental maps of chromatin looping and transcription factor binding enable biological interpretations for observed interactions.^8,9^ The study of gene expression is also directly relevant to complex disease: the vast majority of variants identified in genome-wide association studies are non-protein coding, and thus it is presumed that the disruption of gene regulation is causally involved in the development of many common diseases.^10,11^ In several instances, it has been shown that single nucleotide variants regulate gene expression by altering the function of regulatory elements, and that these altered gene expression profiles result in clinical phenotypes.^12,13^ By better understanding the genetic control of gene expression, we may therefore better understand the genetic architectures underlying complex disease.

Genetic variants associated with gene expression levels – termed expression quantitative trait loci (eQTL) – have been studied extensively in primary human tissue and in cell lines. In many eQTL analyses, a gene-based approached is taken wherein variants within the *cis*-regulatory region for a given gene are tested for association with its expression. Until recently, the number of association tests required to perform a similar genome-wide association test for interactions was not computationally feasible. However, advances in computational power are continually diminishing this barrier and two genome-wide studies of epistasis have identified replicating interactions.^14,15^ The validity of these interactions, however, was questioned when it was demonstrated that through complex linkage disequilibrium (LD) patterns, these putative interactions could tag single variant eQTL.^16^ Notably, all of the interactions identified in those studies were either no longer significant or were strongly attenuated when the effects of *cis-* eQTL were considered. This illustrates that, compared to single-locus analyses, the statistical models used to detect epistasis are subject to novel confounding factors, which are rarely addressed in studies of epistasis.

In this study, we investigate whether evidence for epistasis in humans persists after systematically accounting for technical, statistical, and biological confounding factors. We performed a targeted investigation of interactions regulating gene expression levels in human lymphoblastoid cell lines (LCLs): the analysis was restricted to nominal eQTL within the target gene’s cis-regulatory region, to drastically reduce the number of association tests performed while retaining the genomic regions most likely to harbor pertinent regulatory elements. Few genes showed evidence of epistasis (165 of 11,465 genes tested), although multiple interactions were often detected for the same gene. A total of 1,119 interactions were identified, many of which replicated in an independent dataset (363 of 803 possible). We then investigated confounding factors – technical (variants within probe binding sites, ceiling/floor effect), statistical (missing genotype combinations, population stratification), and biological (haplotype effects, tagging *cis*-eQTL) – that provide alternative, more parsimonious explanations than biological epistasis. Ultimately, each of the interactions identified could be accounted for by an alternative mechanism, suggesting that the majority of statistical interactions identified without accounting for confounding factors are spurious associations. Many of these confounding factors are inherent to the statistical models used, and will therefore generalize to other phenotypes; consequently, the analytic framework of this study will be of use to many future studies of statistical epistasis.

## Subjects and Methods

### Genotyping and gene expression data

The discovery dataset was comprised of individuals ascertained as part of the International HapMap Project, PhaseI*II^17^, which consisted of 210 unrelated individuals with genome-wide genotyping data (Phase I*II, release 24). For each of these individuals, Stranger et al. collected and normalized gene expression levels from immortalized LCLs using the Sentrix Human-6 Expression Bead Chip, v1.^18^ All probes with a HapMap SNP underlying the expression probe were removed from analysis.^18^ We applied a population normalization procedure, described by Veyrieras et al.,^19^ to the gene expression values that such that the expression of each gene within each population followed a normal distribution. This removed population-level differences in gene expression, which enabled us to combine all ethnicities in our analysis. Our replication dataset consists of 232 unrelated individuals from the 1000 Genomes Project (1KG), for whom gene expression in LCLs was available. These individuals had been sequenced at low coverage as part of the 1KG Project;^20^ we used genetic data from phase I, version 3. Stranger et al. also collected and normalized gene expression levels in LCLs for these individuals using Illumina Sentrix Human-6 Expression BeadChip, v2.^21^ We applied the same population normalization procedure ^19^ to these data. Both the discovery and replication dataset are multiethnic; the sample composition by ethnicity is shown in Table 1.

**Table 1.** Dataset Composition by Ethnicity. The number of individuals of each ethnicity (1KG abbreviations) in the discovery and replication analyses.

| Analysis | Total Sample Size | Ethnicity |  |  |  |  |  |  |  |
| --- | --- | --- | --- | --- | --- | --- | --- | --- | --- |
|  |  | CHB | CEU | GIH | JPT | LWK | MXL | MKK | YRI |
| Discovery | 210 | 45 | 60 | - | 45 | - | - | - | 60 |
| Replication | 232 | 34 | - | - | 35 | 80 | 38 | - | 45 |

Two additional replication datasets were used to investigate a promising interaction. The first consisted of 283 European-descent individuals from the Genotype-Tissue Expression (GTEx) Project, for whom gene expression in whole blood was assessed by RNA-sequencing.^22^ Genotype data for these individuals was collected on both the HumanOmni5-Quad Array and the Infinium Exome Chip, and then imputed to 1KG.^22^ The second dataset consisted of brain samples from autopsied European-descent individuals in the Mayo Late Onset Alzhemier’s Disease Consortium.^23^ These individuals were genotyped on the Illumina HumanHap300-Duo Genotyping Beadchip and gene expression was collected using the Illumina Whole-Genome DASL HT BeadChip.^23^ 370 individuals had expression data available from cerebellum, and 385 had expression in the temporal cortex.

### Generating SNP pairs for interaction testing

To generate SNP-pairs for each gene, we first identified all common SNPs within the gene’s *cis*-regulatory region. To be considered common, variants had to have a MAF > 5% when all ethnicities were combined. Based on *cis*-eQTL analyses,^19^ the cis-regulatory region was defined as starting 500 kb upstream of the gene’s start and ending 500 kb downstream of the gene’s stop (including the gene itself); gene boundaries were taken from ENSEMBL. Previously, these variants were individually tested for association with the gene’s expression level in the discovery dataset by Veyrieras et al.^19^ Based on this analysis, we filtered out SNPs whose marginal effects were not nominally associated with gene expression (excluded p > 0.05), under the hypothesis that nominally associated variants may represent weak marginal effects from a true underlying interaction. We then considered all possible SNP-pairs amongst the remaining variants. Once this was done for each gene, over 21 million SNP-pairs were generated for interaction testing.

### Identifying significant interactions

Each SNP pair was tested for interactions significantly associated with the expression of the gene for which it was generated. The following interaction model (Equation 1) was used:

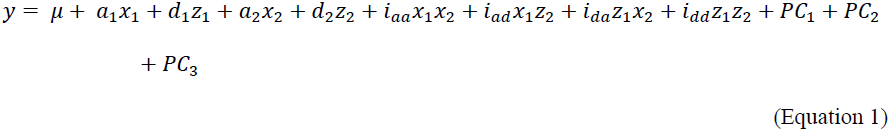

where *y* represents gene expression, *x*_1_ and *x*_2_ use additive encoding to represent the genotype at SNP A and SNP B respectively, *z*_1_ and *z*_2_ use Cordell’s^24^ dominant encoding to represent the genotype at SNP A and B respectively, *a*_1_ and *d*_1_ are estimated coefficients representing the additive and dominant effects of SNP A, *a*_2_ and *d*_2_ are estimated coefficients representing the additive and dominant effects of SNP B, and *i_aa_*, *i_ad_*, *i_da_* and *i_dd_* are estimated coefficients representing both additive and dominant interaction effects. The top three principal components were also included as covariates (*PC*_1–3_). To determine the significance of interactions, this model was compared to a reduced model lacking the four interaction terms using a likelihood ratio test (LRT) (Equation 2).

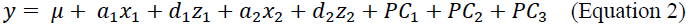

This test was implemented using the program INTERSNP.^25^ We calculated an FDR of 5% using the qvalue package in R^26^

### Identification of representative interaction eQTL models for distinct pairs of interacting genomic loci

Some interaction eQTL (ieQTL) models identified in the discovery analysis were redundant due to LD. For two ieQTL models to be considered redundant, each SNP within one significant ieQTL model had to be in high LD (r^2^ ≥ 0.9) with a SNP within the second ieQTL model, and vice versa. By using this criterion, the pairs were effectively correlated at r^2^ ≥ 0.8, the threshold typically used for tag-SNP selection. The redundant SNP-pairs have very similar betas for all parameters (Supplemental Figure 2), indicating they represent the same signal from a pair of interacting genomic loci. Redundant ieQTL models were grouped together. The model with the most significant LRT p-value in the discovery analysis was used to represent the entire group in most analyses, so that each pair of interacting genomic loci was equally represented. A visual schematic of this process is provided in Figure 1.

**Figure 1.**
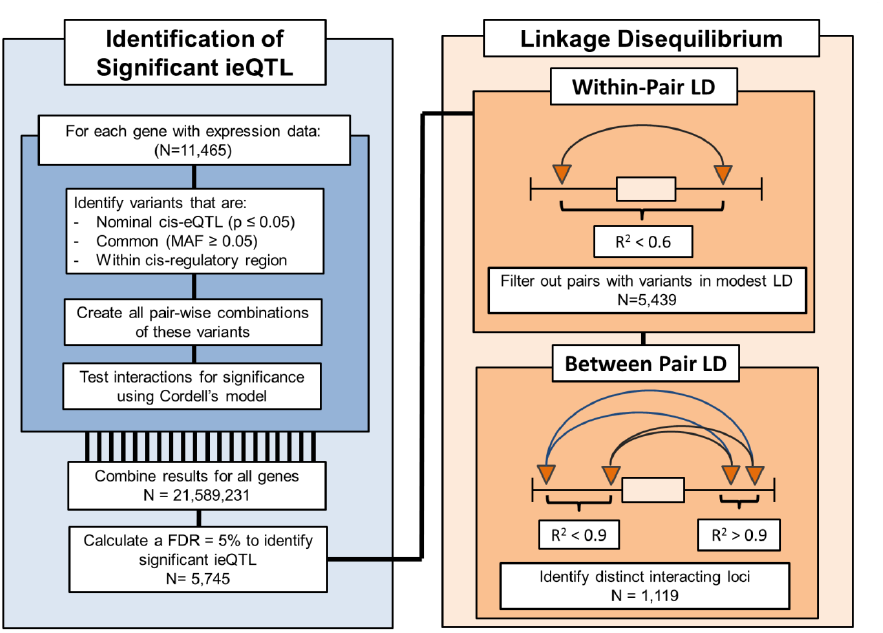
Workflow used to identify and group ieQTL. In the discovery analysis, nominally significant *cis*-eQTL (denoted by triangles) were paired together and tested for interactions significantly associated with gene expression levels (denoted by arcs). The within-pair LD was then calculated (Supplemental Figure 1), and interactions composed of variants in modest LD (r^2^ > 0.6) with one another were removed from the remainder of the analysis. Some of the remaining interactions represented the same pair of interacting genomic loci (Supplemental Figure 2), and were paritioned into distinct groups (denoted by the arc color). For two interactions to be grouped together, each SNP within one significant ieQTL model had to be in high LD (r^2^ ≥ 0.9) with a SNP within the second ieQTL model, and vice versa.

### Variants within the probe binding site

To determine if variants were within the probe binding locations, we first used BLAT to identify the probe binding location in hg19 coordinates. Some probes returned multiple hits; consequently, we filtered the binding sites (binding sites had to be on the same chromosome as the gene, have a length > 30 base pairs, and an identity score > 95%) to identify unique binding locations. We then exclusively looked within a subset of our discovery dataset with sequencing data in the 1KG Project (n=174) to determine if there were any variants within binding sites that might confound the interaction analysis.

### Ceiling/floor effect

Microarrays have a limited dynamic range that is not able to capture the extremes of gene expression. If the combined additive effect of two variants exceeds the threshold of detection, they may be spuriously identified as interacting. We looked for statistical patterns characteristic of a ceiling/floor effect to determine an upper bound of its prevalence within our results. First, we identified the significant (β±SE could not contain zero) variables in the model. All interactions were then categorized as having 0, 1, or 2 SNPs with a significant main effect - either additive or dominant main effects counted; if both additive and dominant main effects were significant for the same variant, the one with the largest effect size was used to represent the main effect. For interactions where both variants had at least one significant main effect, we determined whether or not they had a concordant direction of effect. For those pairs with concordant directions of effect, we compared the significant interaction term with the largest absolute effect size to determine if it was discordant with the main effects. If this was the case, the interaction had a pattern consistent with a ceiling/floor effect.

### Population specific cis-eQTL

Population-specific cis-eQTL can confound the interaction analysis, even though gene expression values were population normalized and the top three PCs were included as covariates. To investigate this, we first stratified the discovery dataset by each of the three ethnicities (CEU, YRI, CHB*JPT), and tested each interaction for significance, using the same methodology. For interactions that were not significant (p < 0.05) in any of the populations, we determined if the interacting variants were population-specific *cis*-eQTL using the following model (Equation 3):

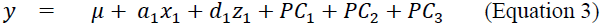

where *y* represents gene expression, *x*_1_ uses additive encoding to represent the genotype for the variant, *z*_1_ uses Cordell’s^24^ dominant encoding to represent the genotype, and the top three principal components were included as covariates (*PC*_*1-3*_). Variants with nominally significant (p < 0.05) main effects were considered *cis*-eQTL. If a variant was identified as a *cis*-eQTL in only a subset of populations, it was considered population-specific.

### Conditional cis-eQTL analysis

To determine if interaction-eQTL pairs were tagging a cis-eQTL as suggested by Wood et al.,^16^ we first identified all nominal cis-eQTL (p < 0.05) for genes with significant ieQTL. To identify all nominal *cis*-eQTL, we used a subset of the discovery analysis individuals (n=174) who were also sequenced as part of the 1KG Project.^20^ We used the called genotypes from Phase III, v5. The same gene expression data previously described for the discovery set was used. Within this subset, we performed a single-marker *cis*-eQTL analysis for each common variant (MAF > 5%) within the *cis*-regulatory region using Equation 4:

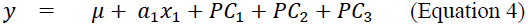

where *y* represents gene expression, *x*_1_ uses additive encoding to represent the genotype for the variant, and the top three principal components were included as covariates (*PC*_*1-3*_). Variants with nominal significant (p < 0.05) main effects were considered *cis*-eQTL.

To determine if any of these *cis*-eQTL could account for the interaction, we created all pairs of *cis*-eQTL and ieQTL for the same gene. We incorporated each *cis*-eQTL into each interaction model (Equation 5) as shown below.

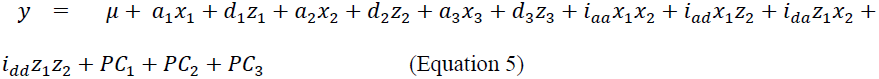

where *y* represents gene expression, *x*_1_ and *x*_2_ use additive encoding to represent the genotype at interacting SNPs A and B respectively, *z*_1_ and *z*_2_ use Cordell’s dominant encoding to represent the genotype at interacting SNPs A and B respectively, *a*_1_ and *d*_1_ are estimated coefficients representing the additive and dominant effects of SNP A, *a*_2_ and *d*_2_ are estimated coefficients representing the additive and dominant effects of SNP B, and *i_aa_*, *i_ad_*, *i_da_* and *i_dd_* are estimated coefficients representing both additive and dominant interaction effects.. The main effect of the *cis*-eQTL is represented with additive encoding by *x*_3_ and with dominant encoding by *z*_3_; the estimated coefficients corresponding to the main effects are *a*_3_ and *d*_3_ respectively. The top three principal components were also included as covariates (*PC*_*1-3*_). We then performed a LRT comparing this model to a reduced model lacking the interaction terms (Equation 6).

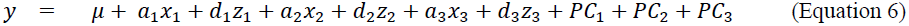

If the LRT p-value of an interaction was nominally significant (p < 0.05) for all conditional analyses, we considered this evidence that the interaction and *cis*-eQTL represented independent signals.

## Results

### Discovery and replication of genetic interactions that impact gene expression levels

We identified interactions between nominal *cis*-eQTL that were significantly associated with gene expression levels. Our analysis was conducted using 210 individuals from the HapMap Project, Phase I*II, on whom both genotyping^17^ and gene expression data within LCLs^18^ were available. A population normalization procedure was applied to the gene expression data, so that there were no systematic differences between populations. The overall workflow for the analysis is shown in Figure 1. For each gene with expression data (n=11,465), we identified common SNPs (global MAF > 5%) within its cis-regulatory region, defined as 500 kb upstream to 500 kb downstream of the gene. To increase power, we only considered variants nominally associated with the gene’s expression (p < 0.05) in a single-marker analysis.^19^ We analyzed all pair-wise combinations of these variants for each gene, resulting in over 21 million SNP pairs. We then performed a likelihood ratio test (LRT) comparing a full model, which contains the top three PCs, main effects, and interaction terms, to a reduced model, containing only the covariates and main effects, to determine which interactions significantly improved model fit.^24^ Given the large number of correlated tests, we controlled the false discovery rate (FDR) at 5% (p ≤ 1.328x10-5) across p-values from all LRT performed.^26^

LD between variants complicates the interpretation of the interaction models. We addressed two types of LD in significant interaction models: within-pair LD, defined as the LD between the variants in the same interaction model, and between-pair LD, defined as the LD between variants in different interaction models. Modest within-pair LD indicates the variants may be identifying a haplotype, which complicates their interpretation because the haplotype may carry other variants that drive the association with gene expression. To protect against confounding by haplotype effects, we removed all pairs with variants in modest LD with one another (r^2^ > 0.6) from the remainder of the analysis. The median r^2^ between remaining pairs of interacting variants was 0.06 (Supplemental Figure 1); however, we examined them for further evidence of haplotype effects (including applying stricter LD filters) in subsequent sections. Ultimately, 5,439 interaction models were both significant and passed the within-pair LD filtering criteria; they were significantly associated with the expression of 165 unique genes (Dataset S1). We then calculated between-pair LD, or the correlation of variants in different interaction models. Highly correlated interaction models were grouped together (Methods, Figure 1) because they likely represent the same pair of interacting genomic loci, as evidenced by their very similar statistical models (Supplemental Figure 2). The 5,439 interaction models represented 1,119 pairs of interacting genomic loci (Dataset S1). The interaction model with the most significant p-value in the discovery analysis was selected to represent the entire group in all subsequent analyses, unless specifically stated otherwise, to ensure that each pair of interacting genomic loci was equally represented.

Next, we performed a replication analysis using an independent dataset of 232 unrelated individuals from the 1KG Project who had both whole-genome sequencing^20^ data and gene expression levels in LCLs^21^ available. All ieQTL composed of variants that were common (MAF > 5%) and had available genotyping data were tested for significant interactions with the same procedure used in the discovery analysis. Of the 803 ieQTL tested, 363 had p-values < 0.05 and 90 passed a Bonferroni multiple testing correction for all tests performed in the replication analysis. We used a liberal threshold, and considered all ieQTL models with LRT p-values < 0.05 as successfully replicated.

### All interactions can be explained by confounding factors

Statistical interactions can be produced by a variety of factors other than biological epistasis, including technical artifacts, statistical artifacts, and LD artifacts driven by other biological processes. Technical artifacts are caused by the limitations of the data itself; for instance, limitations in the dynamic range of measureable gene expression can result in interactions being identified through the ceiling/floor effect. Statistical artifacts are caused by improper application of statistical methodology; for example, when there are population-level differences in the phenotype, analyzing multiple ethnicities together can produce spurious associations due to population stratification. Technical and statistical artifacts are especially troubling since they are unlikely to represent real biological association between the loci and phenotype. Other biological phenomena, namely haplotype effects and *cis*-eQTL effects, can be captured by interaction analyses due to LD patterns. We investigated whether the observed significant ieQTL models could be explained by each of these phenomena.

### Some statistical interactions are consistent with confounding by technical limitations

The gene expression data used in this analysis was collected using microarrays. Microarray technology has a limited dynamic range, meaning that the upper and lower bound on the level of gene expression that microarrays can detect does not cover the full range observed in nature. When the observed range of gene expression values is limited due to technical constraints, variants with sufficiently large main effects may mask the main effects of other variants in the model if their combined effect exceeds the range limitation.^27^ This phenomenon, referred to as the ceiling/floor effect, may result in the identification of spurious interactions. Interactions caused by the ceiling/floor effect have a characteristic pattern, in which the main effects of both variants have the same direction of effect and the interaction terms are in the opposite direction. For example, both main effects may increase gene expression, but the interactions will decrease gene expression. An example of an interaction putatively caused by the ceiling effect is shown in Figure 2. Of 1,119 locus pairs, 48 exhibited a pattern consistent with the ceiling/floor effect. Since transcript production may also have a true biological ceiling, it is possible that true genetic interactions could produce this pattern; consequently, we consider this an upper bound of the influence of ceiling/floor artifacts within our analysis.

**Figure 2.**
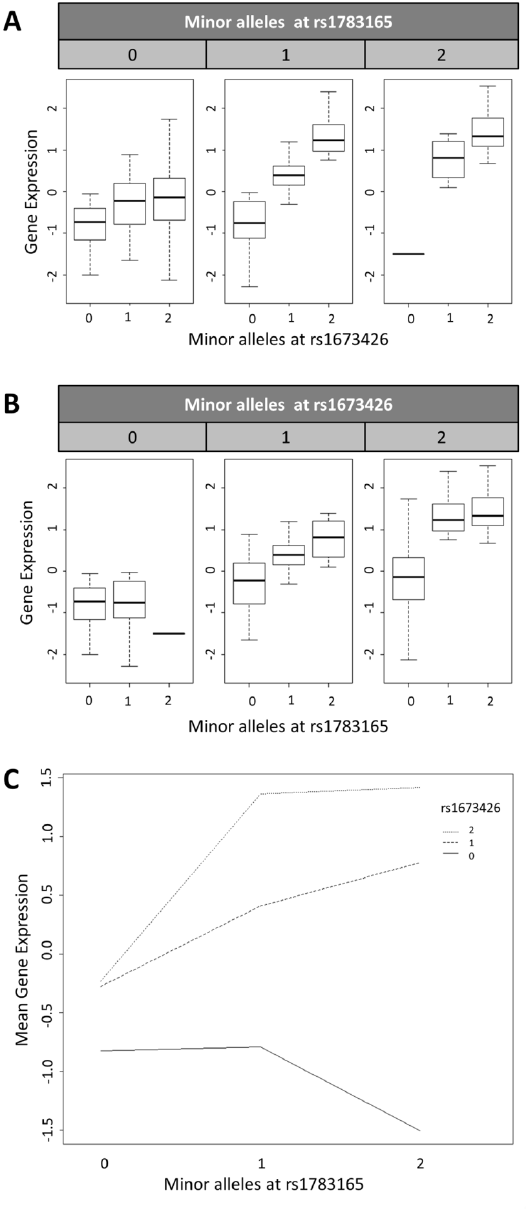
The interaction between rs1783165 and rs1673426 associated with the expression of *PKHD1L1* may be a ceiling effect. The ceiling effect, caused by limitations in the detectable range of gene expression, has a hallmark pattern – both variants have main effects with concordant direction of effect, and the interaction term has a discordant direction. (A) The minor allele of rs1673426 increases the expression of *PKHD1L1*. (B) The minor allele of rs1783165 also increases the expression of *PKHD1L1*, meaning both variants have a concordant direction of effect. The interaction plot (C) depicts the mean gene expression for all individuals with the specified genotype combination. When there is only one minor allele at rs1673426, the mean gene expression increases for each minor allele at rs1783165; however, when there are two minor alleles at rs1673426, the increase in gene expression due to minor alleles at rs1783165 reaches a ‘maximum’ at one minor allele. There is no additional increase in expression for having two minor alleles at rs1783165. This is denoted by the flat line connecting the two genotype combinations. Given that each minor allele at rs1783165 increases gene expression on the background of one minor allele at rs1673426, and that the ‘maximum’ reached on the background of two minor alleles at rs1673426 is very close to the maximum gene expression levels possible to observe, we consider this an example of the ceiling effect.

The interpretation of microarray data is also complicated by genetic variants in the probe binding site, as different alleles may have different affinities for the probe. Probes containing any HapMap variant had previously been removed from the analysis;^18,19^ however, HapMap does not provide comprehensive coverage of genetic variants. Consequently, we looked in a subset of individuals from the discovery analysis (n=174) with low-coverage sequencing data through the 1KG Project to see if genetic variants within the probe binding site may result in apparent interactions. The probes for 508 of 1,119 ieQTL contained a SNPs or indel in the 1KG Project. The probes for 255 ieQTL contained at least one common (MAF > 5%) variant. While the conditional analysis (Methods) would likely account for the effect of these variants, we did not consider ieQTL with a common variant in the binding site evidence for biological epistasis. The probes for the remaining 253 ieQTL contained at least one rare variant, but no common variation. To determine if these rare variants could result in the interaction, we performed the interaction analysis using only the 1KG individuals who did not have a rare variant in the probe binding site. The interactions for 200 ieQTL remained nominally significant (p < 0.05) when all individuals with rare variants were removed. Consequently, the interactions for 811 ieQTL are not attributable to variants within the probe binding sites.

### Missing genotype combinations may result in ieQTL

Linear regression models for epistasis may be unable to accurately decompose variance between genetic terms if there is either LD between the interacting variants or if there are missing genotype combinations. The issue of LD has previously been explored, and the Cordell model is robust to LD between variants when all genotype combinations are present.^28^ Consequently, we examined all interactions within the discovery dataset to see if all of the nine possible two-locus genotype combinations were present. For 457 of the 1,119 ieQTL, at least one genotype combination was absent. While failure to see certain two-locus genotypes may be due to lethal combinations, and thus perhaps is evidence for epistasis, it may also simply be a result of certain combinations being uncommon due to allele frequencies. Either way, the statistical model used cannot provide robust estimates unless all genotype combinations are present, and therefore, we do not consider these interactions to provide evidence for biological epistasis.

### Haplotype effects captured through complex LD patterns may produce ieQTL

In some LD architectures, a combination of two variants can identify haplotypes. While there is evidence to suggest haplotypes form in response to biological interactions between variants,^29^ haplotypes may simply carry other variants that additively regulate gene expression. Figure 3 illustrates how additional variants on the haplotype may result in statistical interactions.

**Figure 3.**
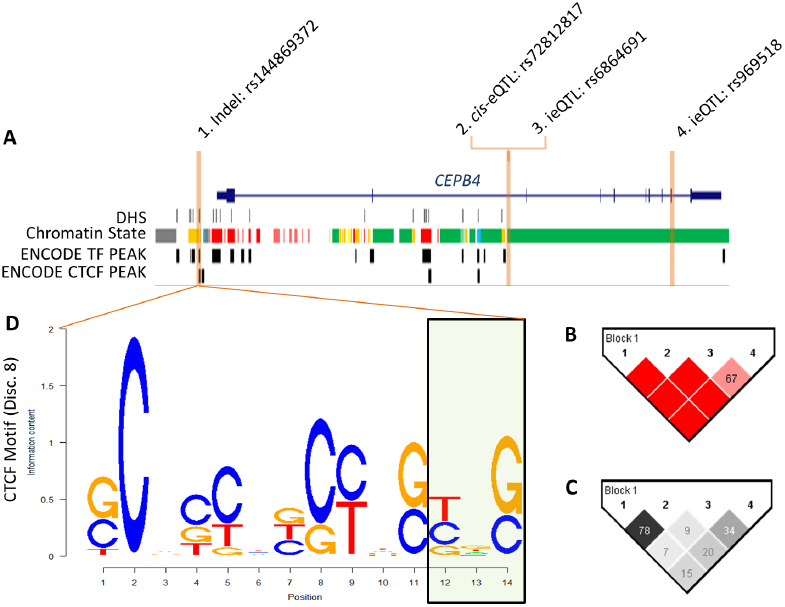
Interactions impacting the expression of *CPEB4* may represent haplotype effects. (A) A significant interaction between rs6864691 and rs969518 regulating the expression of *CPEB4* was identified that replicated and was inconsistent with other confounding factors. The *cis*-eQTL rs72812817 mediated this interaction in the conditional analysis; however, none of these variants were within putative regulatory elements in GM12878 assayed by the ENCODE Project. However, a D' heatmap (B) of the region (the numbers correspond to SNP labels in A) illustrated that an indel, rs144869372, always occurred on the background of the c/s-eQTL (D' = 1). (C) This occurs despite modest r^2^ values, as shown in the r^2^ heatmap of the region. (A) There is evidence from ENCODE suggesting the structural variant may be functional, as it occurs within both a ChromHMM strong enhancer (yellow) and a CTCF binding peak in GM12878. (D) Notably, the structural variant is predicted to alter the binding of CTCF by HaploReg, by altering the last three nucleotides in the binding motif. Given the functional genomics evidence, the indel may be the causal variant, which is detected through interactions that tag the haplotype carrying the indel.

Consequently, interactions between variants on the same haplotype cannot be used to demonstrate the existence of ieQTL in this analysis. As previously stated, we removed all interaction models composed of variants in modest LD with one another (r^2^ > 0.6) from all portions of the study. We additionally investigated whether or not variants within the same interaction model were in modest LD with one another as measured by D’. Of the 1,119 interacting loci, 806 had D’ values < 0.6. The distribution of LD statistics, both r^2^ and D’, for interaction models is shown in Supplemental Figure 1.

### Population specific eQTLs may produce statistical interactions

In our discovery and replication analyses we analyzed multiple ethnicities together. When there are population differences in *both* the distribution of genotypes and phenotypes, analyzing multiple populations together can lead to spurious results. The population normalization procedure applied to the gene expression data removes systematic population differences in the phenotype, thereby enabling multiple ethnicities to be combined for analysis without risk of known complications from population stratification. While this approach has been used in other studies, we also controlled for the top three PCs in our analysis to adjust for residual ethnicity-dependent effects.^19,30^ Furthermore, we performed a stratified analysis, wherein we tested each of the 1,119 ieQTL in each of the three discovery ethnicities (CEU, YRI, and CHB*JPT) separately. While the Cordell model was not robust in the stratified analysis in many cases (due to the reduced sample size, all nine possible two-locus genotype combinations were often not observed in all populations), 859 of 1,119 ieQTL were nominally significant (p < 0.05) in at least one population, suggesting that population stratification is unlikely to account for their significance.

However, the interaction for 260 ieQTL was completely attenuated in the stratified analysis, suggesting that interaction testing was subject to a novel form of confounding by population stratification. Indeed, we found that 234 of 260 ieQTL attenuated in the stratified analysis involved at least one population-specific *cis*-eQTL, meaning that while the variant was present in all populations, it operated as a *cis*-eQTL in only a subset.^21^ The systematic differences between the main effect of each variant and the frequency of two-locus genotype combinations between populations resulted in a spurious interaction signature; an example is provided in Figure 4. To investigate whether population-specific effects may impact the 859 ieQTL that were nominally significant in at least one population, we calculated the within-population LD between each pair of interacting variants. 689 of 859 ieQTL were significant in at least one population where the variants were not in LD with one another (r^2^ and D’ < 0.6) (Dataset S2). We did not consider the 170 ieQTL that were exclusively significant in populations with population-specific haplotypes as evidence for biological epistasis. Ultimately, 689 of the 1,119 ieQTL did not appear to be driven by population-specific effects.

**Figure 4.**
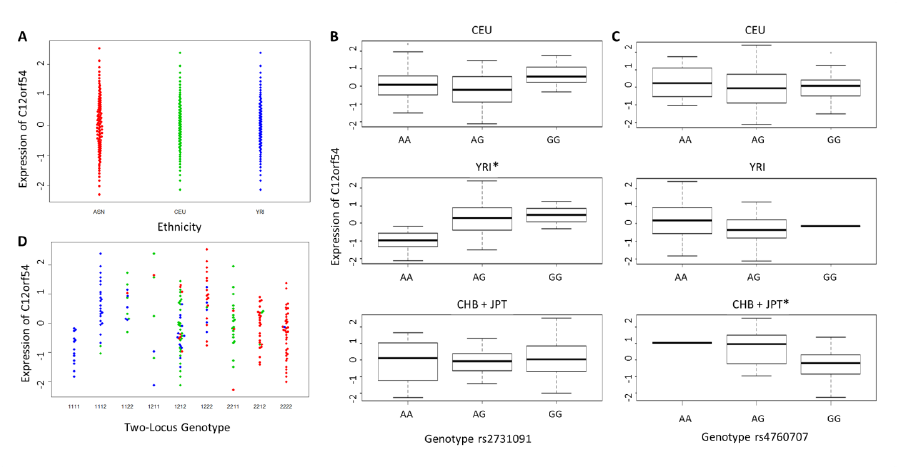
Population specific eQTLs may underlie ieQTL regulating *C12orf54*. The interaction between rs2731091 and rs4760707 regulating *C12orf54* replicated, but was not nominally significant (p < 0.05) in any population in the stratified analysis. (A) Due to the population normalization procedure, there are not systematic differences in the expression of *C12orf54* between populations; however, we found that each variant was a population-specific *cis*-eQTL. (B) rs2731091 significantly regulated gene expression as a *cis*-eQTL in YRI (p = 7.28x10^−6^), but not CEU (p = 0.14) or CHB*JPT (p=0.84). (C) rs4760707 was a cw-eQTL in CHB*JPT (p=7.25x10^−6^), but not in YRI (p=0.17) or CEU (p=0.96). (D) There were clear population differences in the frequency of two-locus genotypes between populations; in combination, it appears the population differences in two-locus genotypes and population specific *cis*-eQTL produced a nuanced form of population stratification.

### Statistical interactions may tag single-variant cis-eQTL through LD patterns

It was recently demonstrated that all the ieQTL identified in one genome-wide association study of epistasis could be explained by the effects of *cis*-eQTL.^16^ To illustrate this, the most significant *cis*-eQTL for each gene with an interaction was identified and the interaction was then conditioned on it (i.e., the *cis*-eQTL was included in the interaction model). When this was done, no interaction remained significant. This occurs when the two interacting SNPs together tag a single *cis*-eQTL through complex LD patterns. Rather than examining only the most significant *cis*-eQTL for each gene, we conditioned all interactions on all nominal *cis*-eQTL (p < 0.05) identified for the regulated gene. This approach requires a comprehensive list of *cis*-eQTL; consequently, we performed this analysis in a subset of individuals from our discovery dataset (n=174) with sequencing data available through the 1KG Project. While the 1KG sequencing data is low coverage, it is extremely unlikely we would fail to detect the effect of a common *cis*-eQTL – 1KG estimates they had 99.3% power to detect variants of 1% frequency.^20^ Even if a common *cis*-eQTL was missed, all variants that could tag it through LD would additionally have to be absent for its effect to not be captured in the conditional analysis. In the 1KG data, all common variants (MAF > 5%) within the *cis*-regulatory region that were nominally associated (p < 0.05) with gene expression were considered *cis*-eQTL. We then created all pairs of *cis*-eQTL and ieQTL for the same gene. For each of these combinations, we performed a conditional analysis in which the additive and dominant main effect for the *cis*-eQTL were incorporated into both the full and reduced model used in the LRT to determine the significance of the interaction. The majority of interactions appeared to be mediated by *cis*-eQTL (Figure 5); however, 139 of the 965 testable ieQTL remained significant (p < 0.05) in all conditional analyses performed, indicating that these interactions are not explained by *cis*-eQTL.

**Figure 5.**
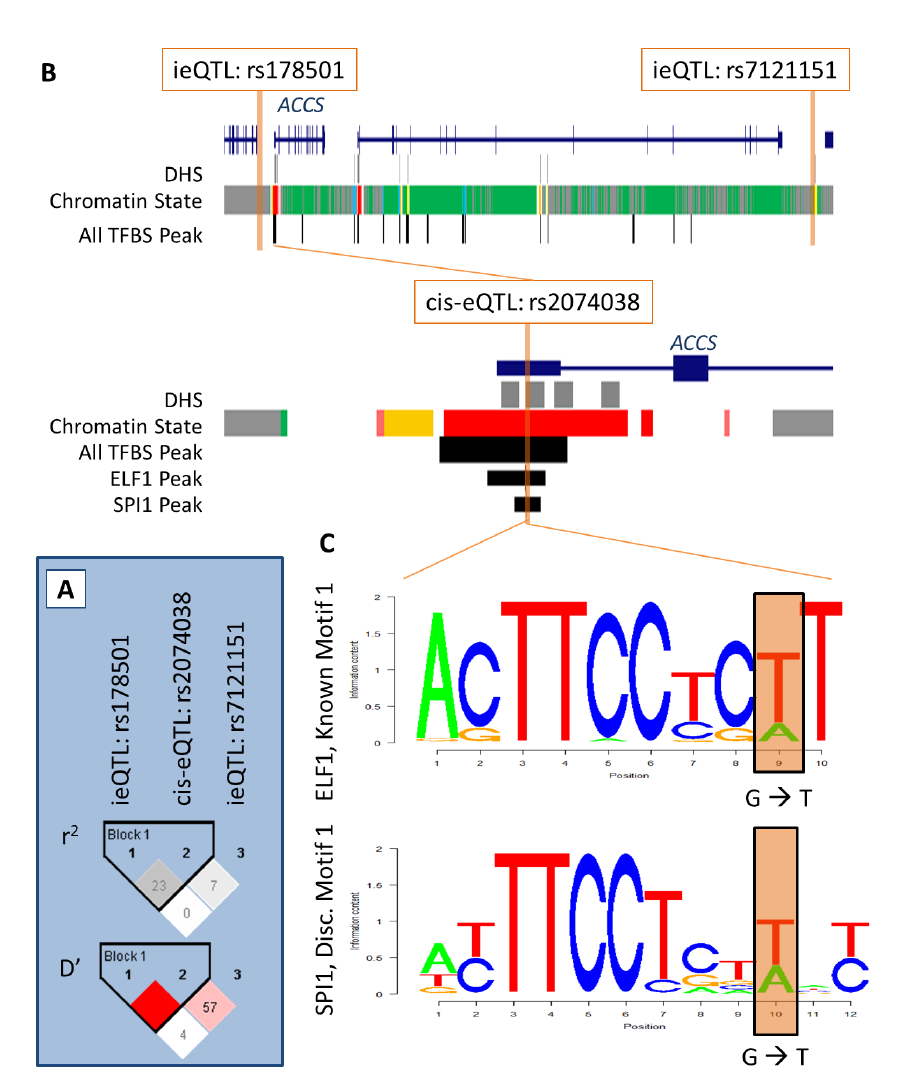
The interacting SNPs regulating *ACCS* are likely tagging a single-variant *cis*-eQTL through linkage disequilibrium. The interaction between rs178501 and rs7121151 is mediated by the c/s-eQTL rs2074038 in the conditional analysis (interaction p-value > 0.05). (A) While the interacting variants are in low LD with the c/s-eQTL based on r^2^, their high D’ indicates they often occur on the same haplotype. (B) The interacting variants are not located within DNase hypersensitivity sites, predicted chromatin states with a regulatory function, or any of the uniform binding peaks identified for all transcription factors tested in GM12878 by ENCODE; however, the *cis*-eQTL is located within the canonical promoter for *ACCS*, a DNase hypersensitivity site, and numerous transcription factor binding peaks identified in GM12878 by ENCODE. (C) Notably, the *cis*-eQTL occurs within a binding peak for both ELF1 and SPI1 in GM12878, and also alters the binding motifs of these transcription factors at the position highlighted in orange. Thus, the *cis*-eQTL rs2074038 is likely the causal variant, and the interaction is simply capturing its effect through LD.

### IeQTL can be entirely accounted for by alternative mechanisms

Finally, we assessed the cumulative impact of alternative explanations on interaction models (Dataset S2). Of the 1,119 interacting genomic loci identified, 363 replicated in an independent dataset. Of these, 68 ieQTL could be explained by technical artifacts (i.e., the ceiling/floor effect and/or variants within the probe binding sites). 199 of the remaining 295 ieQTL could be explained by statistical artifacts (i.e., population stratification and/or missing genotypes).

Biological explanations other than epistasis – namely haplotype effects or the tagging of *cis*-eQTL – could account for 94 of these ieQTL. Ultimately, two interactions (Table 2) replicated and were not explained by the ceiling/floor effect, population stratification, variants within the probe binding site, missing genotype combinations, haplotype effects, or the tagging of *cis*-eQTL.

**Table 2.** Interactions that replicate and are not accounted for by other explanations.

| Gene | SNP 1 | SNP 2 | Discovery‡ | Replication‡ | $r^2$ | D' |
| --- | --- | --- | --- | --- | --- | --- |
| *MYRFL* | rs1262808 | rs11615099 | 7.41 | 2.69 | 0.11 | 0.36 |
| *APIP* | rs1549791 | rs7115749 | 5.00 | 2.54 | 0.12 | 0.43 |

‡ $-\log_{10} P$ values for 4 d.f. LRT interaction test.

We further investigated these two interactions. The interaction between rs1549791 and rs7115749 to regulate *APIP* was not consistent in the direction of effect between the discovery and replication datasets (Supplemental Figure 3). This interaction was only observed in African-descent populations, and without additional datasets to further investigate it, we do not consider it evidence for epistasis due to lack of consistency. The interaction between rs1262808 and rs11615099 regulating the expression of *MYRFL* had concordant effects in both the discovery and replication datasets (Figure 6). As this interaction was observed in European-descent populations, we were able to validate this interaction in 283 individuals from the Genotype-Tissue Expression (GTEx) Project with RNA-sequencing of gene expression in whole blood, and again found a similar pattern of effect (Figure 6). The same trend was also observed in 370 European-descent individuals with gene expression in both cerebellum and temporal cortex, illustrating that the interaction was found in very different cellular conditions and was robust in four independent datasets (Figure 6). However, given that the conditional *cis*-eQTL analysis was conducted in the multi-ethnic discovery dataset, it was possible that population-specific LD patterns could have obfuscated the signal from a single-variant and resulted in the residual significance of the interaction. Consequently, we performed conditional *cis*-eQTL analyses in the additional datasets composed only of European-descent individuals – and found *cis*-eQTL that completely attenuated the significance of the interaction signal in all cases. While the *cis*-eQTL that most attenuates the signal varies between datasets, all tag the same locus (Figure 6). The same locus also attenuates the interaction completely in a conditional analysis on the CEU subset of the discovery dataset (Figure 6). Thus, despite consistent replication in numerous datasets, this interaction can be explained by confounding by *cis*-eQTL.

**Figure 6.**
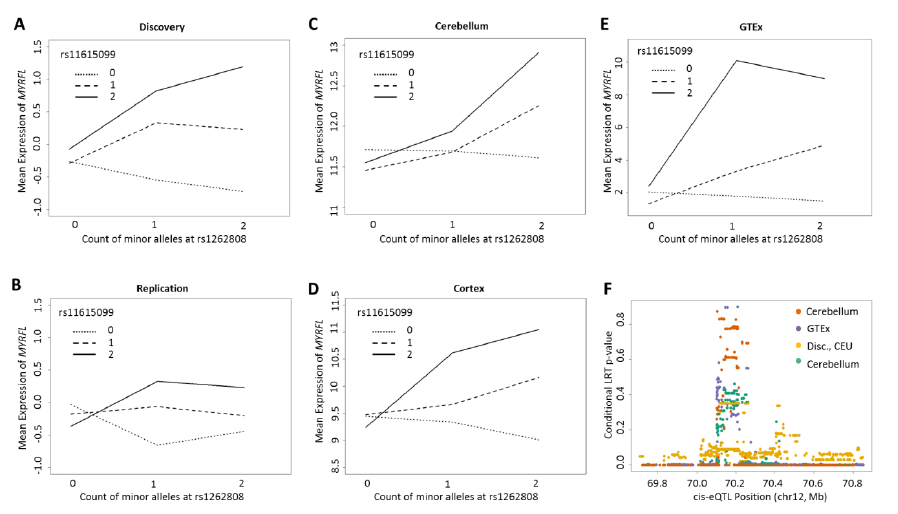
Despite consistent replication, the interaction regulating *MYRFL* is attributable to *cis*-eQTL. In each interaction plot, all individuals are categorized according to their two-locus genotype at rs1262808 and rs11615099. The mean expression of *MYRFL* for all individuals with each of the nine possible two-locus genotypes is shown here for the (A) discovery; (B) replication; (C) Mayo, cerebellum; (D) Mayo, cortex; (E) GTEx, whole blood datasets. The interaction plot illustrates a consistent trend across all datasets, this interaction is mediated by *cis*-eQTL. (F) Conditional *cis*-eQTL analyses were conducted in the discovery (CEU only, yellow); GTEx (purple); Mayo, cerebellum (teal); and Mayo, temporal cortex (orange). For each conditional analysis, the conditional LRT p-value is plotted by the genomic position of the *cis*-eQTL conditioned on. The p-value peak observed in this region illustrates that *cis*-eQTL completely attenuate the interaction when they are conditioned on.

## Discussion

In this study, we demonstrated that many technical, statistical, and biological factors confound the interpretation of statistical interactions identified and replicated in human LCLs. We first detected interactions for less than 2% of genes (165 of 11,465 tested); while this is certainly an underestimate and would increase with sample size, it is markedly lower than the prevalence of *cis*-eQTL.^19^ We then performed a comprehensive investigation of confounding factors (haplotype effects, ceiling/floor effect, single variant eQTL tagged through LD, missing genotype combinations, population stratification, and others), and found that all the interactions identified could potentially be accounted for by at least one of these confounding factors. Consequently, we find no clear evidence for interactions between variants within the *cis*-regulatory region regulating gene expression in humans.

In a range of model organisms, however, epistasis is commonly found^1–3^ – this discrepancy in prevalent genetic architectures may be accounted for by a variety of factors. First, humans are an ‘outbred’ population, whereas model organisms typically are from lab strains with a fairly homogenous genetic background between individuals. If epistatic interactions are actually between numerous variants, the greater diversity in both number of variants and their frequencies observed in outbred populations may prevent any pair-wise interaction from demonstrating a consistent trend at the population level. Additionally, many studies of epistasis in model systems are based upon hybrid crosses, and use linkage to identify broader regions associated with the phenotype of interest. Such studies are more likely to have detected gene-gene or haplotype interactions – which may represent multiple genetic variants at each locus – rather than SNP-SNP interactions. Finally, studies in model organisms that apply similar statistical association designs may also be subject to the confounding factors addressed here.

Confounding factors have a pervasive influence on statistical association studies of epistasis: all the interactions identified in this study could be accounted for by at least one confounder. Our comprehensive exploration of potential confounders, the majority of which are likely to generalize broadly across phenotypes, illustrates the alternative explanations that must be addressed in future studies before concluding that evidence for true biological epistasis has been found. Further, replication – long held as the gold standard for genetic association studies – does not safeguard against these confounding factors, as many are robust to independent replication.

It is also critical to note that other statistical models of epistasis have additional modes of confounding. For example, reduced interaction models that assume exclusively additive main effects for each variant^30–32^ can partition deviations from non-additive main effects (i.e., dominant main effects), into the interaction term. By contrast, the full model used here contains additive and dominant effects for each variant and all four possible interaction terms, thereby allowing variance to be properly partitioned amongst genetic terms.^24^ Additionally, other phenotypes will have distinct forms of confounding; for example, the more removed a phenotype is from the underlying biological process, the more susceptible it is to confounding by threshold effects. To illustrate, imagine that a disease results when the expression of a specific gene exceeds a certain threshold. If that gene has multiple *cis*-eQTL that additively regulate its expression, and only in combination do their effects exceed the threshold, they will be identified as interacting to influence disease risk, even though they behave additively at the mechanistic level. Given the pervasive nature of confounding, it must be considered in all future studies of epistasis. The analytic approach used in this study provides a trait-independent framework for explicitly examining confounding factors and avoiding reporting spurious results.

Future studies of epistasis face many challenges. First, regardless of confounding, epistasis within the *cis*-regulatory region does not appear to be a major component of the genetic architecture underlying the regulation of gene expression in humans. While more statistical interactions with smaller effect sizes would assuredly be identified with increased sample size, the confounding factors would also complicate their interpretation. Furthermore, studies from model organisms have illustrated that epistasis is most commonly observed between variants that exhibit significant single-marker effects^1^; consequently, the study of epistasis is unlikely to identify novel genomic loci or biological pathways related to the phenotype of interest. Instead, studies of epistasis will likely be the most fruitful when looking for modifiers of single-variants that impact a trait of interest, or when trying to determine the cumulative effect (i.e., genetic risk scores) of such variants on the phenotype. Ultimately, the development of high-throughput methodologies to confirm the biological validity of detected interactions at the molecular level will be critical in moving the study of epistasis forward.

## Supplemental Information

**Supplemental Dataset 1. Significant interactions identified in the discovery analysis.** This file provides all 5,439 interactions identified in the discovery analysis. When these interactions appeared to represent the same signal, due to LD, they were placed into groups (n = 1,119) and a representative interaction was chosen. We provide the group identifier for each of the interactions, and the group’s representative interaction.

**Supplemental Dataset 2. Alternative explanations for significant interactions identified in the discovery analysis.** We examined whether or not the 1,119 interactions could be explained by confounding factors. Here, we present which alternative explanations could account for each interaction.

**Supplemental Figure 1.**
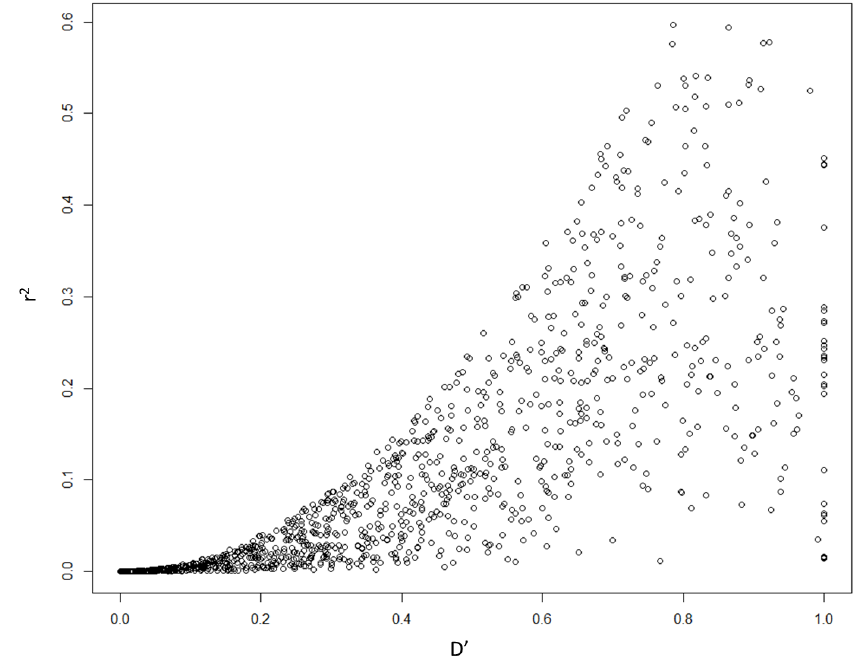
Linkage disequilibrium between interacting variants. We calculated LD between interacting variants using both r^2^ and D’ to determine if they were on the same haplotype. Interactions between variants in modest LD (r^2^ > 0.6) had been removed from all stages of the analysis, and hence are not shown here.

**Supplemental Figure 2.**
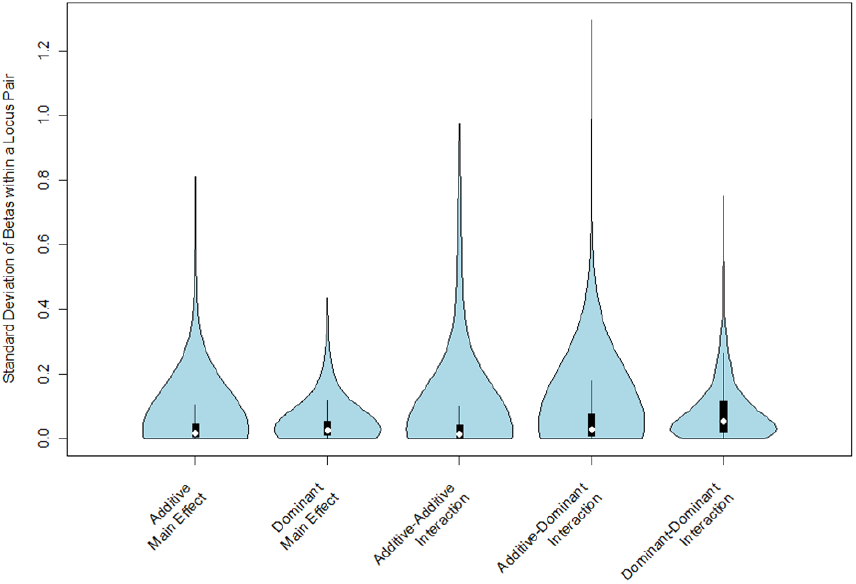
Redundant SNP-pairs have very similar parameter estimates. We grouped together all pairs of interacting SNPs (n=5,439) identified as being redundant through LD measures. For each group, we identified all terms that were significant in at least one of the associated interactions (p < 0.05). We extracted the betas for these significant terms from all interactions within the group. We then calculated the standard deviation of the betas for each significant term within each group to determine how similar the parameter estimates were across all interactions in the same group. The distribution of these standard deviations, categorized by type of variable, is shown above.

**Supplemental Figure 3.**
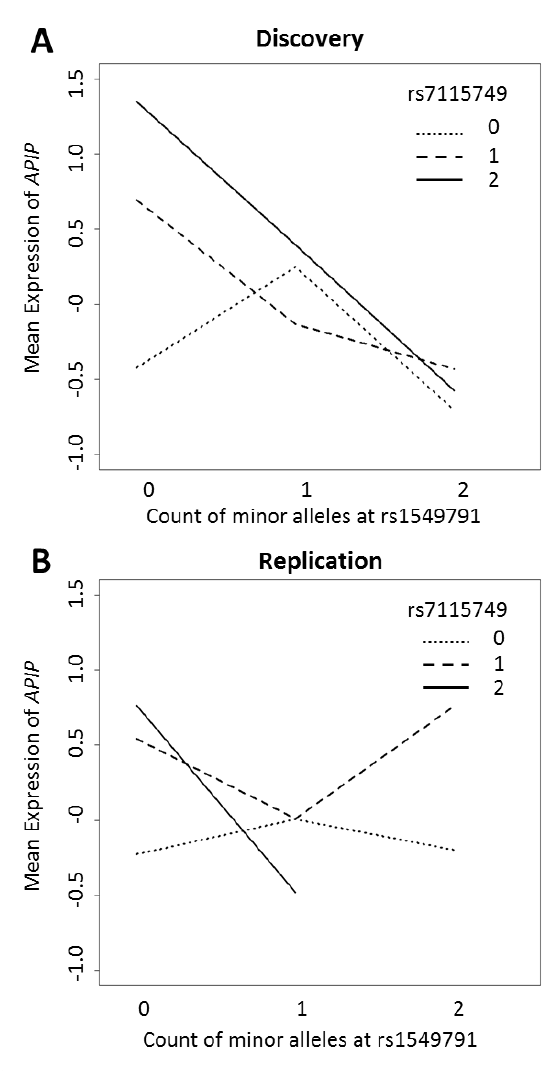
The interaction between rs1549791 and rs7115749 associated with the expression of *APIP* is not consistent between the discovery and replication datasets. In the interaction plot, each individual is categorized according to their two-locus genotype at rs1549791 and rs7115749. This results in nine possible genotype combinations, and the mean expression of *APIP* for each combination is shown here for the (A) discovery and (B) replication datasets. There are markedly different patterns in gene expression by two-locus genotype between the two datasets, illustrating the putative interaction does not replicate with a consistent direction of effect.

## Acknowledgements

We thank Laura Wiley for normalizing gene expression values within the replication dataset. We also thank Jacob Hall, Corinne Simonti, and R. Michael Sivley for their help and advice on this project.

